# Ventral hippocampal interneurons govern extinction and relapse of contextual associations

**DOI:** 10.1101/2023.11.28.568835

**Authors:** Anthony F. Lacagnina, Tri N. Dong, Rasika R. Iyer, Saqib Khan, Mazen K Mohamed, Roger L. Clem

## Abstract

Contextual associations are critical for survival but must be extinguished when new conditions render them nonproductive. By most accounts, extinction forms a new memory that competes with the original association for control over behavior, but the mechanisms underlying this competition remain largely enigmatic. Here we find the retrieval of contextual fear conditioning and extinction yield contrasting patterns of activity in prefrontal cortex and ventral hippocampus. Within ventral CA1, activation of somatostatin-expressing interneurons (SST-INs) occurs preferentially during extinction retrieval and correlates with differences in input synaptic transmission. Optogenetic manipulation of these cells but not parvalbumin interneurons (PV-INs) elicits bidirectional changes in fear expression following extinction, and the ability of SST-INs to gate fear is specific to the context in which extinction was acquired. A similar pattern of results was obtained following reward-based extinction. These data show that ventral hippocampal SST-INs are critical for extinguishing prior associations and thereby gate relapse of both aversive and appetitive responses.

## INTRODUCTION

Animals associate positive and negative events with configural representations of their environments to facilitate the expression of context-appropriate behaviors. However, when the environment changes this learning may no longer be productive or can even become detrimental. Under this circumstance, extinction is a form of long-term memory that attenuates responding in the absence of positive or negative reinforcement^1,2^. Such behavioral flexibility plays a pivotal role in recovery from trauma and addiction, but its efficacy is limited by the tendency for extinguished behavioral responses to re-emerge^3–6^. A potential solution to relapse might entail biasing the expression of extinction memory over previously learned associations.

Contextual learning is probably best understood in fear conditioning, which is thought to be acquired through experience-dependent synaptic plasticity and represented at the neural circuit level by an assembly of hippocampal neurons^7,8^. Extinction attenuates behavioral and neural correlates of fear but does not abolish the original threat association, whose expression can be recovered by certain triggers^9^. A favored explanation has been that fear and extinction memories are stored alongside one another and compete for control over behavior^10–16^, but the mechanisms controlling their selection remain unclear. Indeed, while both forms of learning induce spatial remapping of hippocampal neurons^17–20^, an important unanswered question is whether neurons that gate extinction retrieval are functionally distinct in terms of their molecular or physiological properties from those involved in fear expression. Furthermore, the implications for extinction under normal and pathological conditions will depend on whether such a mechanism is specific to threat-based behavior or generalizes to other forms of learning.

To answer these questions, we performed a Fos-based activity screen of brain regions linked to fear conditioning and extinction. We found that regulation of context fear was associated with contrasting patterns of activity in the prelimbic cortex (PL) and ventral CA1 region of hippocampus (vCA1), where SST-INs were found to be selectively engaged by either fear or extinction retrieval. In vCA1, optogenetic manipulation of SST-INs elicited bidirectional changes in fear expression that were both context- and learning-specific and could not be mimicked by PV-IN stimulation. Similar recruitment of vCA1 SST-INs was observed following extinction of contextual reward conditioning, and suppression of SST-INs evoked relapse of reward seeking. Our results suggest that extinction of contextual associations is controlled by a discrete form of GABAergic inhibition, which could be a general mechanism for discriminating between conflicting internal models of an ambiguous environment.

## RESULTS

### Retrieval of contextual fear extinction preferentially activates vCA1 SST-INs

While expression of fear memory depends on activation of specific cell populations associated with a discrete cue or context, extinction is thought to form a competing association that signals safety in the presence of previously threatening stimuli. Accordingly, successful recall of extinction recruits sparse neuronal populations whose reactivation contributes to low fear expression^12–14,16^. Less understood is whether distinct circuit elements, e.g. neurons with unique molecular or anatomical features, might facilitate extinction memory or promote its expression over conditioned fear. To begin addressing this question, we used c-Fos immunohistochemistry to identify brain regions preferentially activated by extinction retrieval. In addition, given our prior work establishing an important causal role for SST-IN signaling in fear memory encoding^21,22^, we specifically investigated whether activation of region-specific SST-IN populations differs in extinguished versus non-extinguished mice.

To model extinction retrieval, male and female SST-Cre::Ai9 transgenic mice were first submitted to contextual fear conditioning followed by extinction training entailing 5 daily exposures (10 min) to the conditioned context (Ext Ret group, Fig. 1a). Another subset of animals received no extinction training and instead remained in their home cages (Fear Ret group). Additionally, to control for independent effects of repeated context exposure during extinction, a third group was treated identically to Ext Ret mice except that foot shocks were omitted during conditioning (No Shock group). On the final day, all mice were re-exposed to the conditioned context and then sacrificed for immunohistochemical analysis of prefrontal, hippocampal and amygdala subregions, which form a network highly implicated in fear memory^23^. Importantly, freezing during the final test was strongly attenuated in Ext Ret compared to Fear Ret mice, and did not differ from No Shock controls (Fig. 1b).

**Figure 1.**
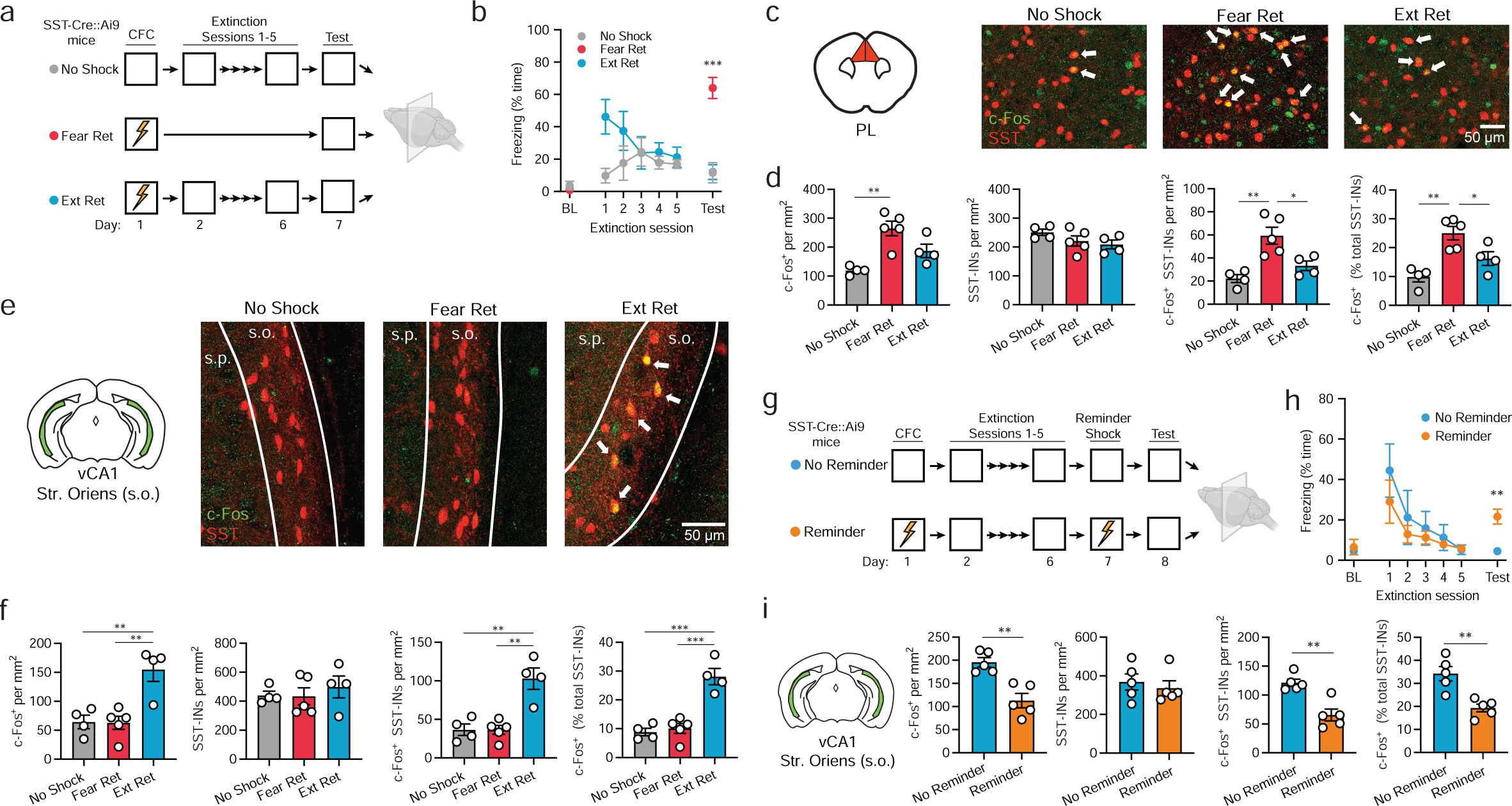
Extinction modulates context-evoked activity of prelimbic and vCA1 somatostatin interneurons. **(a)** Design for analysis of c-Fos expression following retrieval of contextual fear extinction (Ext Ret), as compared to control subjects for which either foot shocks (No Shock) or extinction (Fear Ret) were omitted. Contextual fear conditioning (CFC) consisted of three 2 s 0.75 mA shocks and extinction consisted of 10 min of context exposure daily for 5 days. Brains were collected for immunohistochemical analysis 90 min following a final context test (5 min). All mice were SST-Cre::Ai9. **(b)** Freezing across extinction and test sessions. BL = preconditioning baseline. Freezing during test: *F*_(2, 10)_ = 26.8, *p* < 0.0001, one-way ANOVA. **(c)** c-Fos (red) and SST-IN (green) labeling for prelimbic cortex. Arrows denote co-labeled cells. **(d)** Group comparisons of prelimbic cell counts by one-way ANOVA. c-Fos^+^ cell density: *F* _(2, 10)_ = 11.8, *p* < 0.01. c-Fos^+^ SST-IN density: *F* _(2, 10)_ = 11.4, *p* < 0.01. c-Fos^+^ SST-INs (% total SST-INs): *F* _(2, 10)_ = 12.3, *p* < 0.01. **(e)** c-Fos and SST labeling for vCA1 stratum oriens (s.o.). Arrows denote co-labeled cells. **(f)** Group comparisons of vCA1 s.o. cell counts by one-way ANOVA. c-Fos^+^ cell density: *F* _(2, 10)_ = 11.11, *p* < 0.01. c-Fos^+^ SST-IN density: *F* _(2, 10)_ = 14.17, *p* < 0.01. c-Fos^+^ SST-INs (% total SST-INs): *F* _(2, 10)_ = 27.10, *p* < 0.0001. **(g)** Design for analysis of c-Fos expression following reminder shock-induced relapse. Subjects were treated identically throughout contextual fear conditioning and extinction, except one group was exposed to a single reminder shock prior to test (Reminder). **(h)** Freezing across extinction and test sessions. BL = preconditioning baseline. Freezing during test: *t* _(8)_ = 4.27, *p* < 0.01. **(i)** Group comparisons of vCA1 s.o. immunohistochemical cell counts by unpaired t-test. c-Fos^+^ cell density: *t* _(8)_ = 4.50, *p* < 0.01. c-Fos^+^ SST-IN density: *t* _(8)_ = 4.58, *p* < 0.01. c-Fos^+^ SST-INs (% total SST-INs): *t*(8) = 4.17, *p* < 0.01. **a-f**: No Shock: *n* = 4; Fear Ret: *n* = 5; Ext Ret: *n* = 4. **g-i**: Ext Ret: *n* = 5; Reminder: *n* = 5. Data are presented as mean ± s.e.m. * *p* < 0.05, ** *p* < 0.01, *** *p* < 0.001 by Tukey’s post-hoc test **(b, d, f)** or unpaired t-test **(h, i)**.

Among the regions analyzed, prelimbic cortex exhibited greater c-Fos immunolabeling as well as a higher density and proportion of c-Fos^+^ SST-INs in Fear Ret mice relative to both remaining groups (Fig. 1c-d), suggesting that, similar to prior observations following auditory fear conditioning, contextual fear recall engages prelimbic circuitry involving SST-INs and this activation is reversed with extinction learning^21,22^. However, no differences were noted in either the total number of c-Fos^+^ cells or c-Fos^+^ SST-INs in infralimbic cortex, lateral or basal amygdala (Supplementary Fig. 1a, c-d). Likewise, a similar number of immunopositive cells was observed in the granule cell layer of the dentate gyrus and stratum pyramidale (s.p.) of area CA1 and CA3, in both the dorsal and ventral hippocampus (Supplementary Fig. 1b, e-n). However, c-Fos was elevated in Ext Ret mice within vCA1 and to a lesser extent dorsal CA1 in stratum oriens (s.o.) (Fig. 1e-f; Supplementary Fig. 1j), which is populated primarily by GABAergic neurons and contains the majority of hippocampal SST-INs^24^. Moreover, a higher density and overall proportion of vCA1 SST-INs were found to be co-labeled for c-Fos in Ext Ret mice. Thus, recall of fear and extinction memory elicits contrasting patterns of activity in prefrontal cortex and vCA1, where SST-INs are preferentially activated following extinction. Curiously, there was additional modulation of c-Fos expression within dorsal dentate hilar SST-INs (Supplementary Fig. 1f), but we concentrated our efforts in this study to understanding the functional role of vCA1 populations.

One possibility is that vCA1 SST-INs become engaged following extinction to facilitate the expression of newly adapted behavior (i.e. cessation of freezing). Alternatively, increased vCA1 SST-IN activity may reflect changes driven by extinction learning but not necessarily related to extinction retrieval. Importantly, extinction learning is susceptible to relapse, in which expression of the original fear association returns despite successful extinction learning^9^. We therefore examined c-Fos expression under a relapse scenario to further test modulation of SST-IN activity by high and low fear states. As above, SST-Cre::Ai9 mice received contextual fear conditioning followed by extinction training (Fig. 1g). One day later, they were returned to the conditioned context, where one group of mice received a single reminder shock (Reminder group). This condition is a well-established trigger of relapse, potentially mediated by relearning, reinstatement, or some combination thereof^9^. During a final test session, the Reminder group exhibited higher freezing levels in the conditioned context compared to animals for which the reminder shock was omitted (Ext Ret group, Fig. 1h). Correspondingly, animals receiving the reminder shock displayed fewer c-Fos^+^ cells in s.o. as well as co-labeled SST-INs in vCA1 (Fig. 1i). This further indicates that SST-IN activation is associated with low fear expression and subject to opposing modulation by extinction and relapse.

During our analysis we noted that similar to other recent studies^25,26^, SST-Cre::Ai9 mice exhibit conditional expression of tdTomato in a subset of neurons in s.p. with pyramidal morphology in vCA1 and vCA3 (Supplementary Fig. 2a). Immunolabeling revealed the vast majority of tdTomato^+^ cells in s.o. but not the s.p. express somatostatin (Supplementary Fig. 2e-g). However, to determine whether more selective targeting of SST-INs can be achieved, we compared these results to those from SST-FlpO::Ai65 mice, in which reporter expression relies on Flp rather than Cre recombination. As in the SST-Cre line, SST-FlpO::Ai65 mice exhibited a similar abundance of tdTomato expression in s.o. and tdTomato^+^ cells were predominantly immunopositive for SST. However, in contrast to SST-Cre::Ai9 mice, SST-Flp::Ai65 mice were relatively lacking in pyramidal cell labeling (Supplementary Fig. 2). We therefore selected the SST-FlpO line for subsequent *in vivo* manipulations of SST-INs to minimize potential off-target effects.

### Extinction is associated with alterations in SST-IN synaptic drive

Prior reports indicate that SST-INs in dorsal CA1 (dCA1) express long-term potentiation at excitatory synapses and exhibit experience-dependent plasticity following contextual fear conditioning^27,28^. We therefore wondered whether vCA1 SST-INs might, at a more remote timepoint after conditioning, exhibit either fear or extinction-related alterations in synaptic transmission. SST-Cre::Ai9 mice were submitted to fear conditioning with or without subsequent extinction using the same behavioral design as in Fig. 1, except a final test of memory retrieval was omitted and instead we prepared transverse acute slices of vCA1 (Fig. 2a). Importantly, extinction yielded a reduction in freezing across training sessions (Extinction group, Supplementary Fig. 3). SST-INs were targeted for whole-cell recordings based on tdTomato expression and were further differentiated from pyramidal neurons by their cellular morphology and passive membrane properties. For each recorded cell, spontaneous excitatory (sEPSCs) as well as inhibitory postsynaptic currents (sIPSCs) were analyzed.

**Figure 2.**
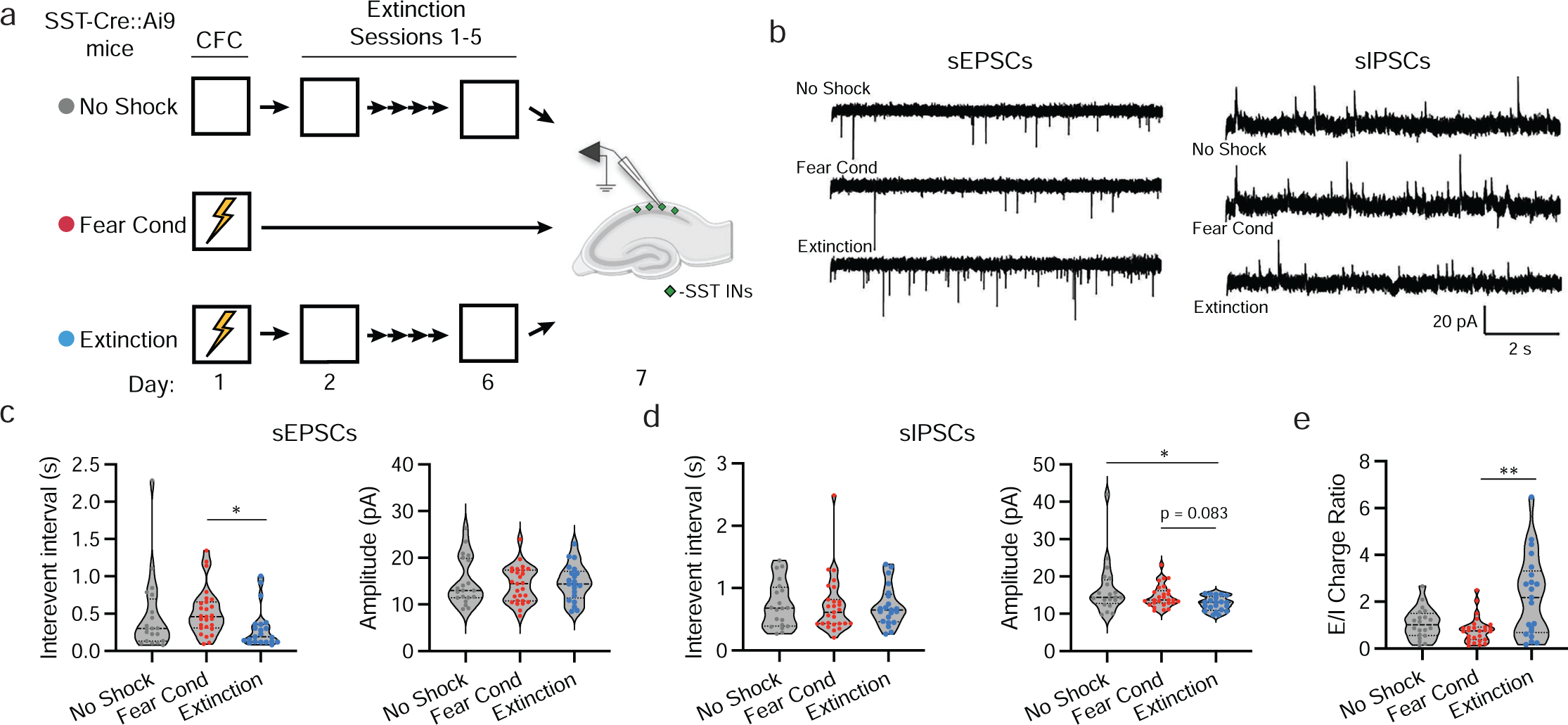
vCA1 SST-INs exhibit altered synaptic transmission following context fear extinction. **(a)** Design for analysis of synaptic transmission using whole-cell electrophysiology following extinction (Extinction), as compared to control subjects for which foot shocks (No Shock) or extinction (Fear Cond) were omitted. All mice were SST-Cre::Ai9. **(b)** Representative current traces containing spontaneous EPSCs (sEPSCs) and sIPSCs, which were sampled from the same population of SST-INs. **(c)** Group comparison of sEPSC properties by Kruskal-Wallis ANOVA. Interevent interval: χ^2^ = 8.14, *p* < 0.05. **(d)** Group comparison of sIPSC properties by Kruskal-Wallis ANOVA. Amplitude: χ^2^ = 7.91, *p* < 0.05. **(e)** Group comparison sEPSC charge normalized to sIPSC charge for individual SST-INs (E/I Ratio) by Kruskal-Wallis ANOVA. Charge (Q) is calculated as the total area of spontaneous currents per second of recording. E/I Ratio: χ^2^ = 10.4, *p* < 0.01. No Shock: *n* = 20 cells (6 mice); Fear Cond: *n* = 27 (4 mice); Extinction: *n* = 21 cells (5 mice). Data are presented as mean ± s.e.m. * *p* < 0.05, ** *p* < 0.01 by Dunn’s post-hoc test.

Results indicated that SST-INs from extinguished mice exhibited a higher frequency of sEPSCs relative to non-extinguished mice (Extinction vs. Fear Cond, Fig. 2b-c). However, neither group differed from No Shock controls. Conversely, extinction was associated with lower amplitude of sIPSCs relative to No Shock controls, as well as a statistical trend toward lower amplitude relative to Fear Cond mice (Fig. 2b, d). Given these mixed effects, we examined whether there were any differences in the balance of excitatory:inhibitory transmission at the level of individual SST-INs, relying on a charge-based metric that accounts for both frequency and amplitude of currents (Fig. 2e). This analysis revealed that, compared to SST-INs from Fear Cond mice, SST-INs from Extinction mice exhibited a higher E:I ratio of spontaneous transmission and might therefore be more readily excitable.

### SST-INs control context-evoked freezing after extinction

The above results indicate that high SST-IN activity is inversely correlated with context fear, suggesting that SST-INs trigger or otherwise support extinction recall. We therefore turned to an optogenetic approach in SST-FlpO mice to test whether vCA1 SST-INs modulate freezing, using a within-animal behavioral design in which halorhodopsin (NpHR)-mediated photoinhibition or channelrhodopsin (ChR2)-mediated photoexcitation was conducted at key timepoints following acquisition, extinction and relapse of contextual fear conditioning (Fig. 3a-c, Supplementary Fig. 4). In addition, to establish context specificity of any effects, photostimulation was repeated in an arena distinct from the one where conditioning occurred. Finally, for each manipulation, an opsin-negative control vector (eYFP only) was infused into a separate group of animals to control for independent effects of laser light.

**Figure 3.**
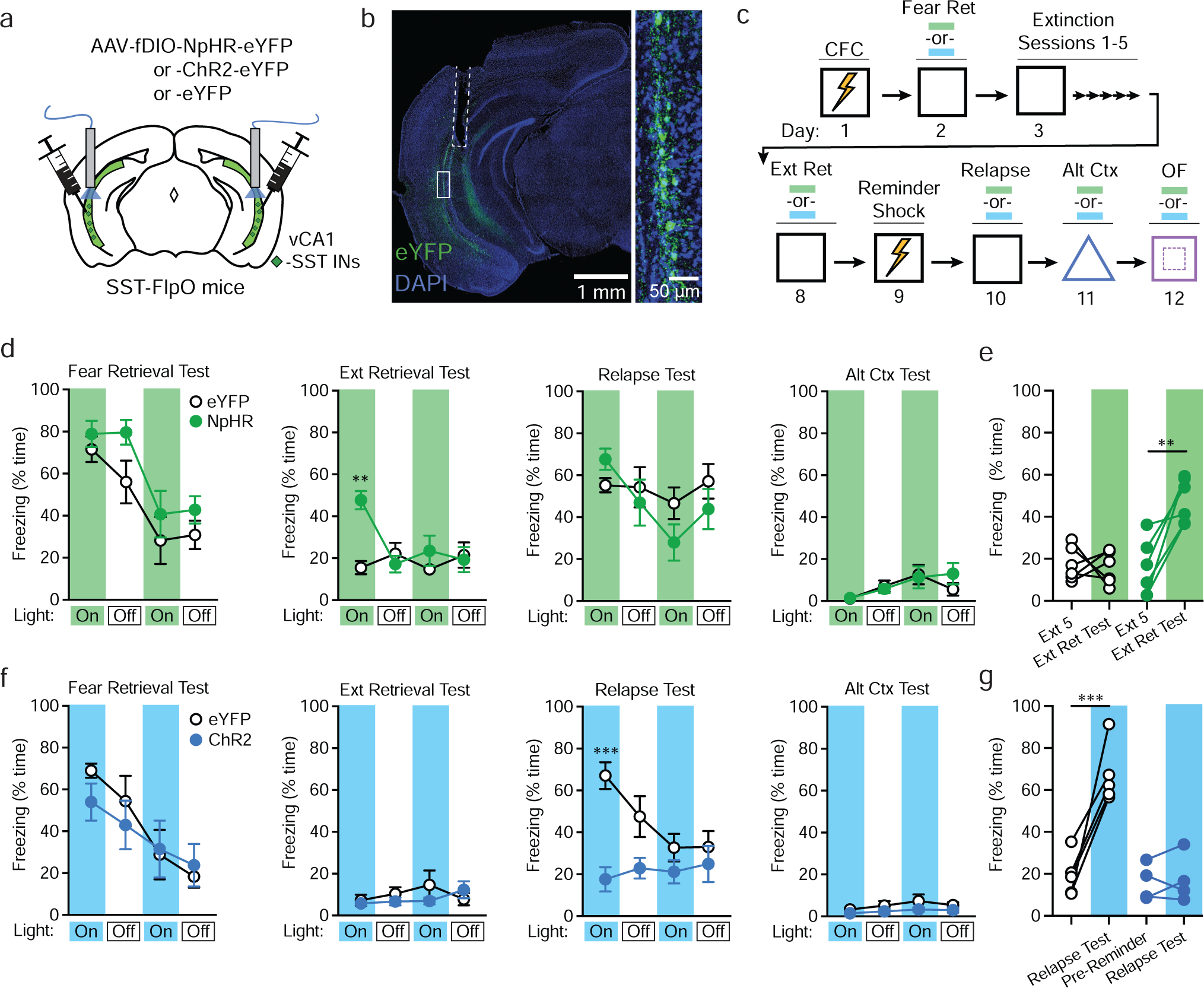
Manipulation of SST-IN activity alters freezing to the conditioned context following extinction. **(a)** Strategy for targeting of vCA1 SST-INs for optogenetic manipulations with halorhodopsin (NpHR), channelrhodopsin-2 (ChR2) or opsin-negative control vectors (eYFP only) in SST-FlpO mice. **(b)** Representative image of resulting viral expression and optic fiber placement (dashed white line). Solid box indicates area of magnification (right). **(c)** Design for contextual fear conditioning (CFC), extinction and reminder shock-induced relapse, with photostimulation tests at the indicated timepoints (blue and green bars). OF = open field test. **(d)** Effect of NpHR-mediated photoinhibition during fear retrieval, extinction retrieval, relapse, and alternate context tests, as analyzed by two-way repeated measures ANOVA across all light-on and light-off epochs (3 min duration). Opsin x epoch interaction: *F* _(15, 150)_ = 2.40, *p* < 0.01. **(e)** Change in freezing between the first 3 min of the final extinction session (Ext 5) and the first light-on epoch of the extinction retrieval test (Ext Ret Test), as analyzed by two-way repeated measures ANOVA. Opsin x test interaction: *F* _(1, 10)_ = 11.1, *p* < 0.01. **(f)** Effect of ChR2-mediated photoexcitation during similar tests as **d**, as analyzed by two-way repeated measures ANOVA across all light-on and light-off epochs. Opsin x epoch interaction: *F* _(15, 105)_ = 2.75, *p* < 0.01. **(g)** Change in freezing between pre-reminder shock baseline period and first light-on epoch of relapse test. Opsin x test interaction: *F* _(1, 7)_ = 20.6, *p* < 0.01. eYFP for NpHR experiment: *n* = 6; NpHR: *n* = 6; eYFP for ChR2 experiment: *n* = 5; ChR2: *n* = 4. Data are presented as mean ± s.e.m. ** *p* < 0.01, *** *p* < 0.001 by Tukey’s post-hoc test **(d, f)** or Šídák’s post-hoc test **(e, g)**.

Prior work indicates that freezing is typically strongest during the initial few minutes of exposure to a conditioned context^13^. We therefore targeted the initial 3 min epoch of each 12 min test for photostimulation, with the rationale that our manipulations would be most likely to influence behavior at this stage. However, a second period of photostimulation was also included to examine the impact of SST-IN activity later in each test session. Analysis of light-on (532 nm, 12-15 mW, constant) and light-off epochs across all tests in NpHR and eYFP mice revealed an opsin x epoch interaction, which could be attributed to higher freezing in NpHR relative to eYFP mice during the extinction retrieval test (Fig. 3d). This effect was specific to the first epoch of the extinction retrieval test, mirroring a prior study involving dentate granule cell inactivation^13^, and did not occur during photoinhibition in an alternate context (Fig. 3d). Further analysis of within-animal freezing (restricted to the initial 3 min epoch) confirmed that NpHR but not eYFP mice exhibited relapse of freezing between the final extinction session and the subsequent extinction retrieval test (Fig. 3d).

We next conducted a similar analysis of behavior during SST-IN photoexcitation (473 nm, 9-12 mW, 20 Hz, 10 ms pulses), which revealed an opsin x epoch interaction across test sessions, as well as lower freezing in ChR2 relative to eYFP mice during the first epoch of the relapse test (Fig. 3e). Further analysis of within-animal freezing confirmed that while eYFP mice exhibited relapse after a reminder shock, this effect was abolished in ChR2 mice (Fig. 3e). However, photoexcitation did not attenuate freezing during the initial test of fear retrieval (Fig. 3e), suggesting that extinction learning was required before SST-INs could exert control over fear expression. Furthermore, there was no effect of photoexcitation in an alternate context, indicating that SST-IN activation did not interfere generally with contextual discrimination processes, leading to fear generalization. Finally, neither NpHR nor ChR2 mice exhibited general locomotor effects upon photostimulation in the open field assay (Supplementary Fig. 4b,d). Thus, vCA1 SST-INs modulate fear under specific scenarios when memories of threat and safety are in conflict. In these situations, SST-IN activity is important for extinction recall and attenuates freezing.

### Control of context fear extinction is a function of specific interneurons

While activating SST-INs reduces freezing under relapse conditions, it is nevertheless possible that stimulation of any large inhibitory population has the potential to disrupt in a broad or nonselective manner the function of vCA1 neurons and may not be indicative of the specific functional capacity of extinction-related SST-INs. Therefore, we first asked whether relapse of extinguished fear can be prevented by reactivating SST-INs that were specifically recruited by extinction learning. For this purpose, we employed a viral genetic strategy entailing expression of an estrogen-receptor-dependent Cre recombinase (ERCreER) under the control of the enhanced synaptic activity responsive element (E-SARE) (Fig. 4a-b)^29^. In a prior report, we demonstrated that this approach effectively labels memory-related SST-INs when used in combination with SST-FlpO driver mice for intersectional recombination^21^. To determine whether E-SARE is suitable for tagging extinction-related SST-INs, we infused AAV-E-SARE-ERCreER into vCA1 along with an INTRSECT eYFP vector, which requires both Cre and Flp recombination for eYFP transcription (Fig 4b). Subjects were then divided into two groups, both of which underwent contextual fear conditioning and extinction, but received injections of 4-hydroxytamoxifen (4-OHT) at different time points to compare resulting cellular tagging (Fig. 4c). Consistent with c-Fos analysis (Fig. 1f), mice that received 4-OHT during the final extinction session exhibited more abundant labeling within vCA1 s.o. than those injected following fear conditioning (Fig. 4d-e). However, labeling within s.p. was exceedingly sparse and not modulated by timing of injection (Fig. 4e), consistent with preferential targeting of SST-INs.

**Figure 4.**
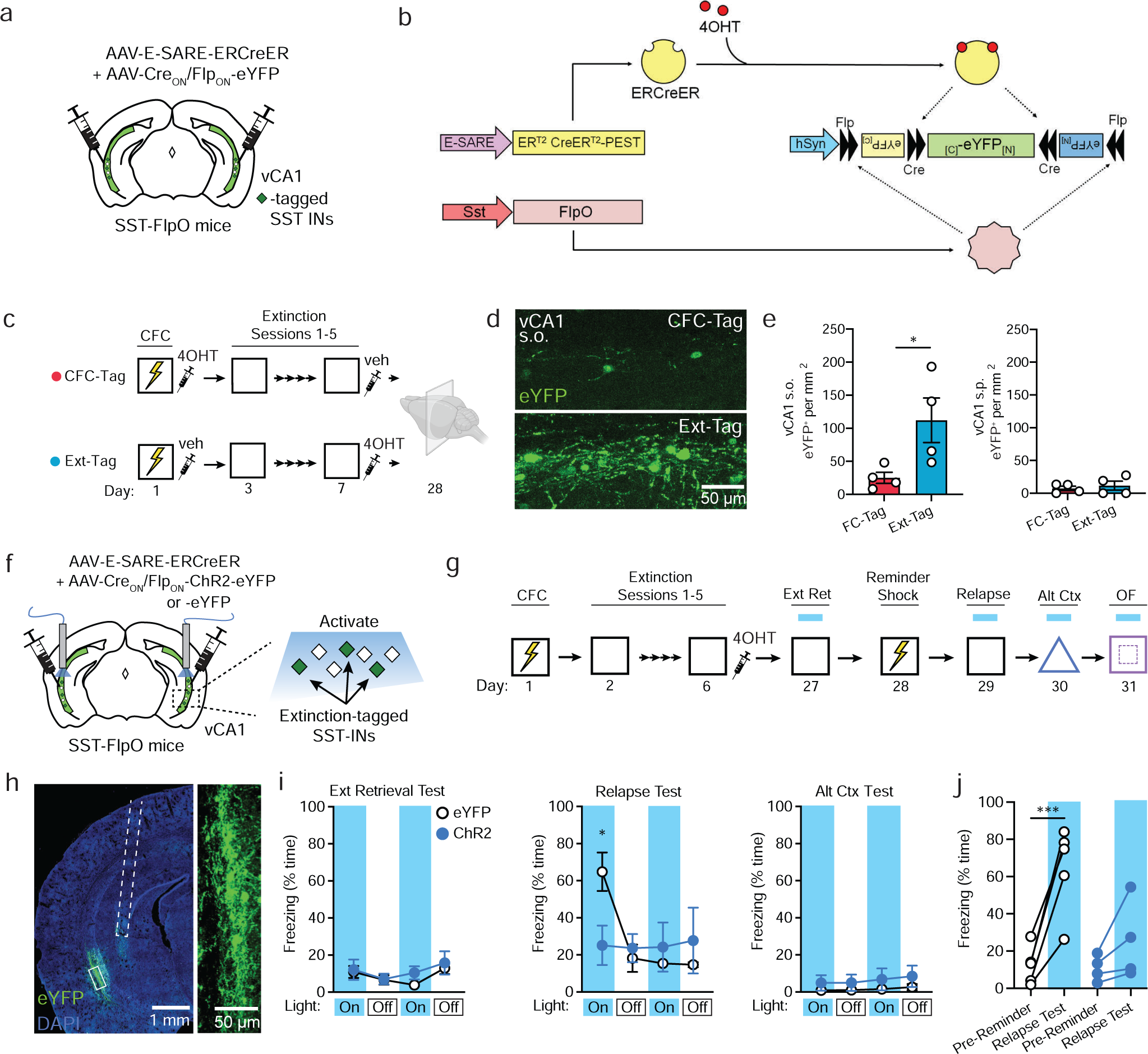
Selective reactivation of extinction-related SST-INs is sufficient to prevent the return of fear. **(a)** Strategy for selective targeting of vCA1 extinction-related SST-INs for expression of eYFP. (**b**) Infusion of viral cocktail into SST-FlpO mice enables intersectional tagging of recently activated SST-INs, temporally restricted by systemic 4-OHT injection. **(c)** Design for SST-IN tagging coinciding with contextual fear conditioning (FC-Tag) or late extinction (Ext-Tag), in which mice were immediately injected with 4-OHT following either session. **(d)** Representative images of eYFP tagging in stratum oriens (s.o.) of vCA1. (**e**) eYFP^+^ cell density within s.o. (left) or stratum pyramidale (s.p., right). vCA1 s.o. eYFP^+^ cell density: *t* _(6)_ = 2.51, *p* < 0.05. **(f)** Strategy for targeting of extinction-related vCA1 SST-INs for optogenetic manipulations with channelrhodopsin-2 (ChR2) or opsin-negative control vectors (eYFP only) in SST-FlpO mice. **(g)** Design for fear conditioning, extinction and reminder shock-induced relapse, with photostimulation tests at the indicated timepoints. OF = open field test. **(h)** Representative image of resulting viral expression and optic fiber placement (dashed white line). Solid box indicates area of magnification (right). **(i)** Effect of ChR2-mediated photoexcitation during extinction retrieval, relapse, and alternate context tests, as analyzed by two-way repeated measures ANOVA across all light-on and light-off epochs (3 min duration). Opsin x epoch interaction: *F* _(11, 77)_ = 3.44, p < 0.001. **(j)** Change in freezing between pre-reminder shock baseline period and first light-on epoch of relapse test. Opsin x test interaction: *F* _(1, 7)_ = 11.7, p < 0.05. **a-e**: FC-Tag: *n* = 5; Ext-Tag: *n* = 4. **f-j**: eYFP: *n* = 5; ChR2: *n* = 4. Data are presented as mean ± s.e.m. * *p* < 0.05, *** *p* < 0.001 by unpaired t-test **(e),** Tukey’s post-hoc test **(i)** or Šídák’s post-hoc test **(j).**

Using the above intersectional genetic strategy, we next targeted extinction-related SST-INs for cellular tagging with either ChR2 or eYFP (Fig. 4f). Following infusion of E-SARE and INTRSECT constructs into vCA1, all subjects underwent fear conditioning and extinction and were injected following the final extinction session with 4-OHT to induce recombination in extinction-related SST-INs (Fig. 4g). Following viral expression, effects of photoexcitation on conditioned freezing were tested during extinction retrieval, relapse, and alternate context exposure. Analysis of light-on and light-off epochs across all tests revealed an opsin x epoch interaction. As with bulk excitation of SST-INs (Fig. 3e), ChR2-expressing mice exhibited lower freezing than eYFP mice during the relapse test (Fig. 4i), which was attributable to a lack of fear return in this group following a reminder shock. No differences were observed during the other context tests or during free exploration of the open field (Fig. 4i, Supplementary Fig. 4f). Therefore, reactivation of a sparse population of extinction-related interneurons is sufficient to attenuate freezing despite their comprising a minority (∼25%) of vCA1 SST-INs.

Nevertheless, while targeting a subset of memory-related SST-INs can avoid potentially nonselective effects, it is nevertheless unclear whether the capacity to inhibit freezing is unique to SST-INs, which are distinct in terms of morphology and synaptic connectivity from other interneuron classes. Parvalbumin-expressing cells (PV-INs) represent the largest subclass of interneurons in vCA1 aside from SST-INs and exert profound control over the firing of excitatory projection neurons^30^. We therefore tested whether extinction-related behavior could be modulated by photoexcitation of PV-INs using a design identical to that employed for SST-IN manipulations (Fig. 3, 5c). In these experiments, PV-Cre mice were employed to restrict expression of Cre-dependent ChR2 or eYFP vectors to PV-INs (Fig. 5a-b). Unlike manipulations of SST-INs, our analysis revealed no effect of opsin expression in these experiments, despite successful acquisition of extinction by both ChR2 and eYFP groups (Supplementary Fig. 4g). Furthermore, both ChR2 and eYFP groups exhibited return of freezing following a reminder shock (Fig. 5d), indicating that PV-IN activation failed to prevent relapse. Thus, fear-attenuating effects of SST-INs cannot be mimicked by other sources of inhibition and are therefore likely to depend on discrete microcircuit dynamics rather than non-selective impairments in hippocampal processing.

**Figure 5.**
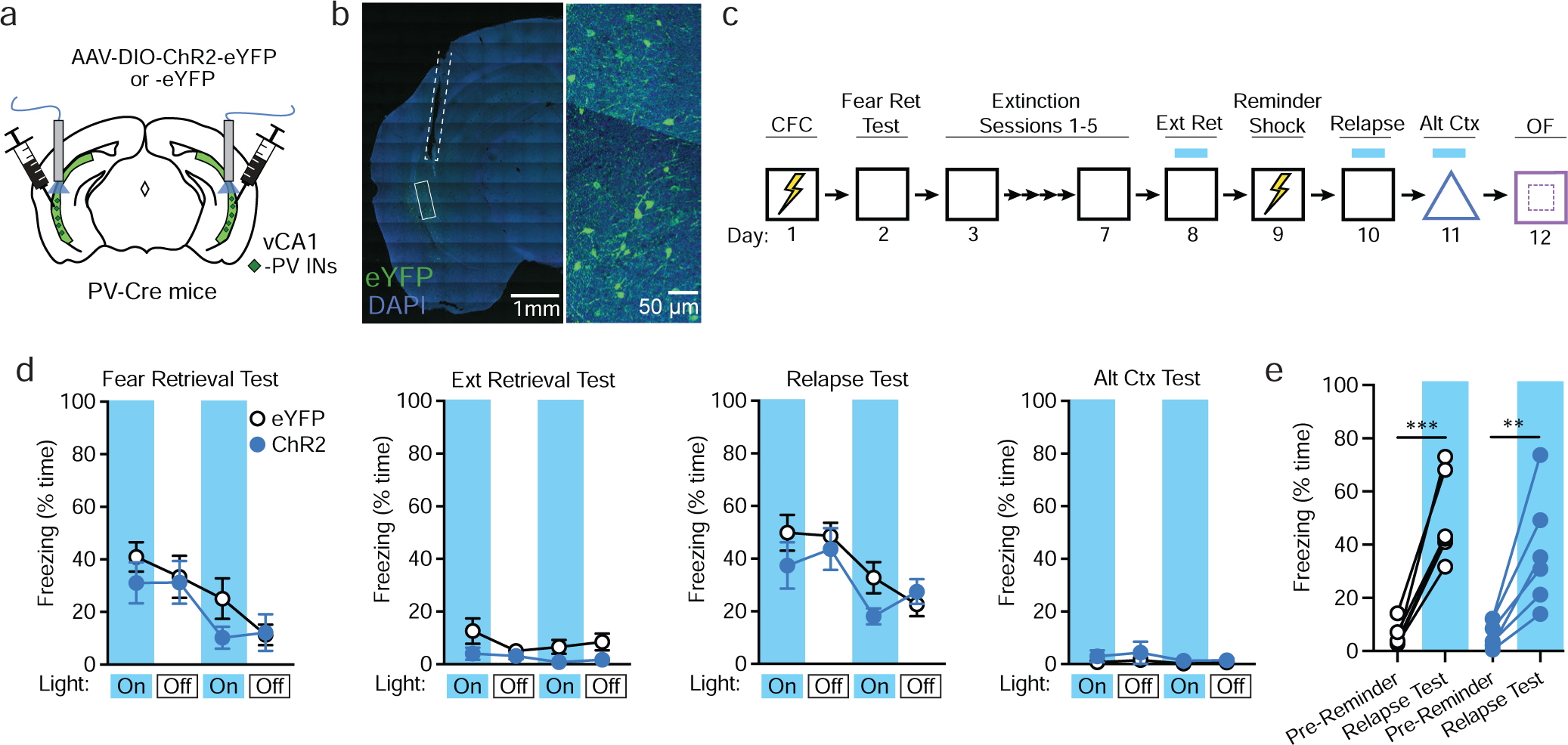
Context fear is unaffected by activation of vCA1 PV-INs. **(a)** Strategy for targeting of vCA1 PV-INs for photoexcitation with channelrhodopsin-2 (ChR2) or opsin-negative control vectors (eYFP only) in PV-Cre mice. **(b)** Representative image of viral expression and optic fiber placement (dashed white line). Solid box indicates area of magnification (right). **(c)** Design for contextual fear conditioning, extinction and reminder shock-induced relapse, with photoexcitation tests at the indicated timepoints. OF = open field test. **(d)** No effect of ChR2-mediated photoexcitation during fear retrieval, extinction retrieval, relapse, and alternate context tests as analyzed by two-way repeated measures ANOVA across all light-on and light-off epochs. **(e)** Change in freezing between pre-reminder shock baseline period and first light-on epoch of relapse test was not modulated by opsin expression. eYFP: *n* = 6; ChR2: *n* = 6. Data are presented as mean ± s.e.m. ** *p* < 0.01, *** *p* < 0.001 by Šídák’s post-hoc test.

### SST-INs also mediate extinction of appetitive contextual associations

The pattern of results obtained following contextual fear conditioning are intriguing in that SST-IN manipulations affected specific stages of behavior. We found no evidence that SST-INs gate freezing prior to extinction learning or in a distinct context where learning did not occur, suggesting that SST-INs are not directly involved in aversive responding but indirectly modulate this behavior by influencing contextual representations. Therefore, to test whether SST-INs support extinction under non-aversive conditions, we implemented a form of appetitive conditioning in which learning is indicated by approach to a reward delivery port. First, adopting a design similar to that used for c-Fos analysis after fear extinction (Fig. 1), we tested whether extinction of contextual reward conditioning activates SST-INs. SST-Cre::Ai9 mice were deprived to 80-90% of free-feeding weight and divided into three groups that were continuously rewarded throughout 14 days of training (Reward Ret), rewarded only on the first 8 days (Ext Ret), or repeatedly exposed to the conditioning context without ever receiving sucrose within the context (No Reward) (Fig. 6a). Reward Ret and Ext Ret groups exhibited increased port entries across training sessions until day 9, after which responding remained high in the Reward Ret condition but was strongly attenuated in Ext Ret mice (Fig. 6b). Following extinction, all groups were returned to the conditioning context and their behavior was examined in the absence of any sucrose delivery before they were sacrificed for immunohistochemical analysis.

**Figure 6.**
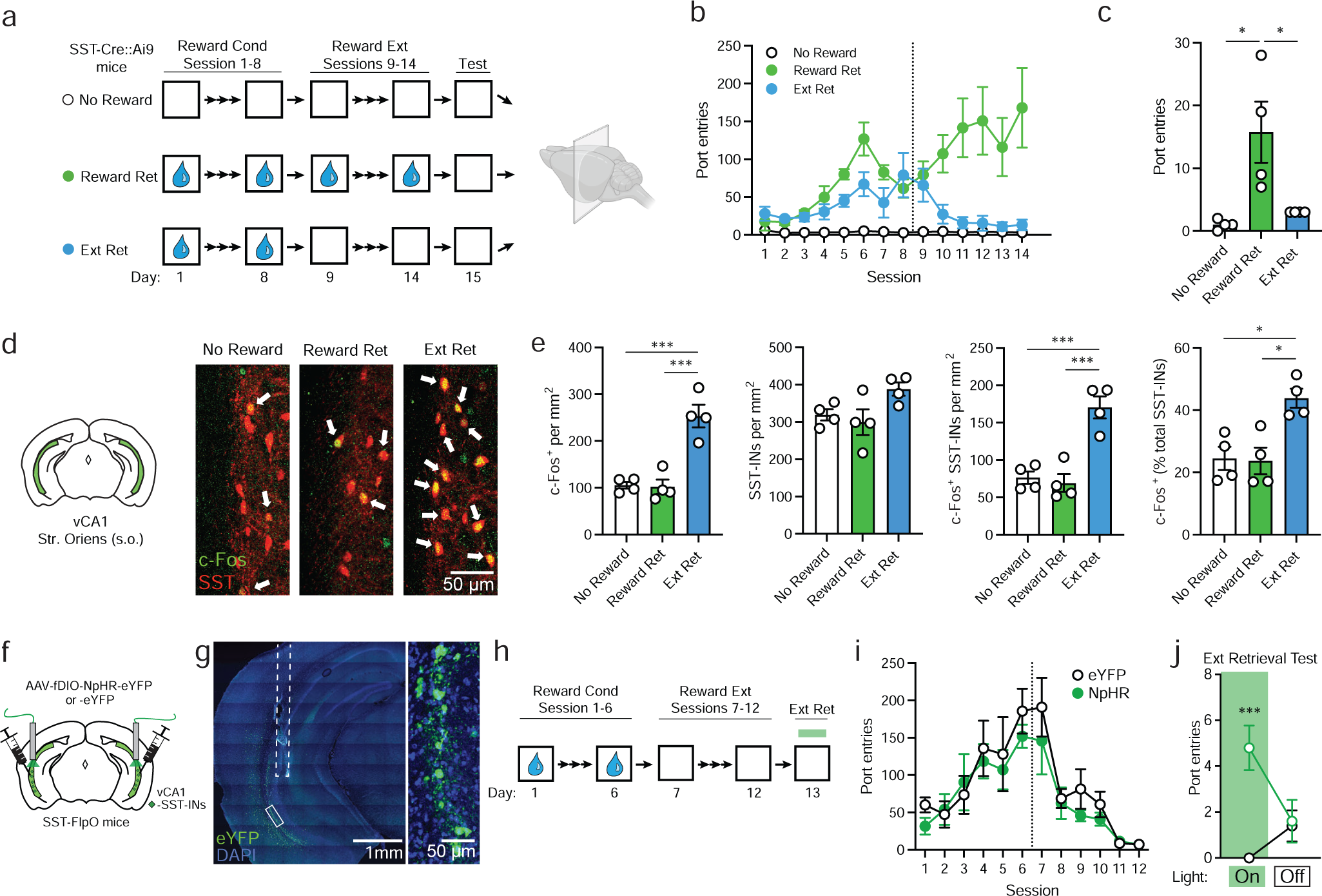
Extinction retrieval after appetitive conditioning depends on vCA1 SST-IN activity. **(a)** Design for analysis of c-Fos expression following retrieval of contextual reward extinction (Ext Ret), as compared to control subjects for which either sucrose reward (No Reward) or extinction (Reward Ret) were omitted. Appetitive conditioning consisted of unsignaled delivery of 30% sucrose to a reward port 30 times a session at a variable interval. Brains were collected for immunohistochemical analysis 90 min following a final context test (5 min). All mice were SST-Cre::Ai9. **(b)** Group comparison of entries into the sucrose delivery port as a function of conditioning and extinction. **(c)** Port entries during 5 min context test where sucrose was omitted in all groups. Port entries during test: χ^2^ = 10.2, *p* < 0.001, Kruskal-Wallis ANOVA. **(d)** c-Fos (red) and SST (green) labeling in vCA1. Arrows indicate co-labeled cells. **(e)** Group comparisons of vCA1 s.o. cell counts by one-way ANOVA. c-Fos^+^ cell density: *F* _(2, 9)_ = 25.8, *p* < 0.001. c-Fos^+^/SST-IN density: *F* _(2, 9)_ = 22.4, *p* < 0.001. c-Fos^+^ SST-INs (% total SST-INs): *F* _(2, 9)_ = 9.40, *p* < 0.01. **(f)** Strategy for targeting of vCA1 SST-INs for photoinhibition with halorhodopsin (NpHR) or opsin-negative control vectors (eYFP only) in SST-FlpO mice. **(g)** Representative image of viral expression and optic fiber placement (dashed white line). Solid box indicates area of magnification (right). **(h)** Design for conditioning and extinction of appetitive responding, with photoinhibition test at the indicated timepoint. **(i)** Group comparison of entries into the sucrose delivery port as a function of conditioning and extinction. **(j)** Effect of SST-IN photoinhibition on sucrose port entries during a 10 min extinction retrieval test, which was divided into 5 min light-on and light-off epochs, analyzed by two-way repeated measures ANOVA. Opsin x epoch interaction during extinction retrieval: *F* _(1, 8)_ = 19.2, *p* < 0.01. **a-e**: No Reward: n = 4; Reward Ret: *n* = 4; Ext Ret: *n* = 4. **f-j**: eYFP: *n* =5; NpHR: *n* = 5. Data are presented as mean ± s.e.m. * *p* < 0.05, ** *p* < 0.01, *** *p* < 0.001 by Dunn’s post-hoc test **(c),** Tukey’s post-hoc test **(e)**, or Šídák’s post-hoc test **(j)**.

Compared to the Reward Ret group, Ext Ret mice made fewer port entries during the final test, and their performance was equivalent to the No Reward group (Fig. 6c). Examination of prefrontal brain sections revealed a greater density of c-Fos positive cells in the prelimbic region of Reward Ret compared to No Reward mice (Supplementary Fig. 5), mirroring group differences observed following contextual fear conditioning (Fig. 1). Likewise, Ext Ret mice exhibited a higher density of c-Fos labeling in vCA1 s.o. and a higher proportion of c-Fos^+^ SST-INs (Fig. 6d-e).

Given these results, we tested whether SST-IN activity influences recall of sucrose reward extinction. SST-FlpO mice received vCA1 injections of Flp-dependent NpHR or eYFP vectors and, following a period of incubation, were submitted to contextual reward conditioning and extinction (Fig. 6f-h). Importantly, both groups exhibited increased port entries as a function of training and an equivalent reduction in responding during extinction (Fig. 6h). Following extinction, animals were returned to the reward context and the behavioral effect of photoinhibition was tested in the absence of reward. Remarkably, NpHR-expressing mice made a greater number of port entries during the test than eYFP mice, consistent with relapse of an extinguished reward association (Fig 6i). This indicates that SST-INs gate extinction retrieval independent of the affective valence of conditioning or whether learning is expressed through an active or passive response.

### Discussion

In this study, we uncovered a neural activity signature of contextual fear extinction retrieval in a genetically discrete population and established a functional role for ventral hippocampal SST-INs in behavioral flexibility. Paralleling extinction-dependent engagement of SST-INs, effects of manipulating this population were highly selective to the context where extinction learning occurred and did not affect initial fear expression. This implies that vCA1 SST-INs are not hardwired to control aversive behavior but rather modulate expression of conflicting contextual responses. Accordingly, our examination of reward-based learning revealed a similar pattern of results, and despite its culmination in an active behavioral output (i.e. approach), we found a shared requirement for SST-INs in the expression of appetitive extinction memory.

The mechanism by which extinction-related SST-INs influence memory processing remains unclear but is likely to depend on their interaction with memory-related excitatory projection neurons. One possibility is that SST-INs suppress the reactivation of projection neurons underlying context conditioning. If true, this is unlikely to result from general network inhibition because we did not observe differences in overall activation of the pyramidal cell layer after extinction (Supplementary Fig. 1b), and behavioral relapse could not be prevented by stimulation of a different inhibitory population (Fig. 5). Instead, our results suggest that the unique synaptic configuration of SST-INs may be relatively more important than their inhibitory output, in which case SST-IN microcircuits exhibit several unique properties that may warrant consideration. First, a large proportion of SST-INs in CA1 preferentially innervate the distal dendritic compartment of pyramidal neurons, where they influence the integration of direct entorhinal inputs^31,32^. Interestingly, layer III neurons of the medial entorhinal cortex, which give rise to these inputs, play an instructive role in reward-based spatial remapping and drive dendritic plateau potentials near reward locations^33^. It is possible that fear conditioning relies on an analogous dendritic signal and that its disruption by SST-IN-mediated inhibition prevents reactivation of a fear-related assembly or, alternatively, stabilizes an extinction-related representation against interference from this type of activity.

Aside from excitatory neurons, SST-INs also synapse extensively onto other interneuron subtypes, including PV-INs and Schaffer-collateral-associated interneurons (SC-INs), for which cholecystokinin (CCK) is a specific marker^34–36^. By interacting with these populations, SST-INs could influence network dynamics through disinhibition to support the recruitment of new cellular assemblies established by extinction learning. While our data appear to exclude PV-INs as potential mediators of such effects, SC-INs are an important source of feedforward inhibition in the Schaffer collateral pathway, where interaction between SST-INs and SC-INs facilitates synaptic integration of CA3 inputs by CA1 pyramidal cells^31^. Thus, signaling from both major afferent pathways controlling CA1 activity can be shaped by SST-INs, potentially in a coordinated fashion that favors processing of some inputs (i.e. CA3) over others (i.e. entorhinal). As the dentate gyrus and CA3 are thought to perform a pattern separation and completion transformation^37^, CA1 SST-INs may play a role in gating information arriving from CA3 in order to disambiguate conflicting memories associated with the context. In addition to pathway-specific control, however, SST-INs support type II theta oscillations, which may contribute to the expression of prior learning through entrainment of discrete learning-related ensembles across hippocampal-prefrontal networks^38,39^.

Just as the downstream effects of SST-IN activity remain to be established, another important question is by what mechanism are they activated in extinction retrieval. Because SST-INs derive a large proportion of their input from local excitatory neurons^24^, it seems plausible that dynamic changes in vCA1 population activity are responsible for their recruitment. In this scenario, SST-INs may participate as a component of an extinction-related assembly whose function is to resolve a conflict between new and prior learning through feedback control of competing populations^40^. Alternatively, inputs originating outside the hippocampus may be more critical in orchestrating the activity and/or plasticity of SST-INs. For example, SST-Ins exhibit potent activation in response to cholinergic signaling^32,41–43^, which may explain a selective requirement for medial septum in extinction but not acquisition of contextual fear^19^.

Regardless of the underlying circuitry, an intriguing facet to our results is that SST-INs exert similar control over aversive and appetitive associations. Interestingly, recent work on olfactory-based conditioning suggests that, in contrast to dCA1, vCA1 preferentially encodes stimuli with emotional significance, and once formed these representations are relatively stable when the conditioned stimulus is paired with an outcome of opposite affective valence ^44^. This has been interpreted to mean that, while vCA1 populations multiplex information about stimulus outcomes, they may also track the current value or salience of a stimulus independent of its associative valence. Future work will be needed to resolve the role of distinct efferent pathways of vCA1 in controlling valence-specific action and whether projections that signal stimulus salience and/or outcome are preferentially modulated by SST-INs.

To conclude, our results suggest that SST-INs are an important substrate for contextual fear extinction that may hold clues about the flexible representation of both aversive and appetitive experiences. When activated, SST-INs function like a mnemonic gate that controls whether new or old learning prevails. As such, they can override established stimulus responses to allow animals to engage in competing adaptive behaviors and potentially encode new stimulus relationships.

## ACKNOWLEDGEMENTS

These experiments were supported by the following grants from the National Institute of Mental Health to R.L.C.: R01 MH116445, R01 MH124880 and R01 MH132224; and a graduate research fellowship from the National Science Foundation to T.N.D. Icons used in Fig. 1a, g, Fig. 2a, Fig. 4c and Fig. 6a were made with BioRender.com. We thank members of the Clem lab for feedback on the manuscript.

## CONTRIBUTIONS

A.F.L. and R.L.C. initiated the project. R.L.C. supervised the research. R.L.C., A.F.L. and T.N.D. designed experiments. A.F.L., T.N.D., R.R.I., S.K., and M.K.M. performed the research and analyzed the data. A.F.L. and R.L.C. wrote the manuscript.

## METHODS

### Animals

Adult male and female mice aged 2-6 months were used in all experiments. All transgenic mice were originally acquired from The Jackson Laboratory and were subsequently bred in-house. Mice were housed in groups of 2-5 in plastic cages with corncob bedding and cotton nestlets and were maintained on a 12 h light-dark cycle (7:00-19:00) in a temperature- and humidity-controlled vivarium. Food and water were provided *ad libitium*, except for appetitive conditioning experiments where food was restricted. All experiments were performed during the light cycle. Mice were randomly assigned to groups prior to experimentation. Sample sizes were chosen to meet or exceed those used in previously published studies using similar experimental designs. The following transgenic mouse lines were used: SST-Cre (stock number 028864; B6J.Cg-*Sst^tm2.1(cre)Zjh^/*MwarJ), SST-FlpO (stock number 028579; *Sst^tm3.1(flpo)Zjh^/*J), Ai9 (stock number 007909, B6.Cg-*Gt(ROSA)26Sor^tm9(CAG-tdTomato)Hze^*/J) mice, Ai65 (stock number 032864; B6.Cg-*Gt(ROSA)26Sor^tm65.2(CAG–tdTomato)Hze^*/J), and PV-Cre (stock number 017320; B6.129P2-*Pvalb^tm1(cre)Arbr^*/J). All experimental procedures were approved by the Institutional Animal Care and Use Committee at the Icahn School of Medicine at Mount Sinai.

### Viral Vectors

Viral vectors purchased from Addgene include AAV1-Ef1a-fDIO-EYFP (#55641), AAV8-nEF-Coff/Fon-ChR2(ET/TC)-EYFP (#137141), AAV8-nEF-Coff/Fon-NpHR3.3-EYFP (#137154), AAV8-hSyn-Con/Fon-EYFP (#55650), AAV8-hSyn-Con/Fon hChR2(H134R)-EYFP (#55645), and AAV1-hSyn-hChR2(H134R)-EYFP (#26973). The plasmid for E-SARE-ERCreER-PEST was obtained as a gift from Dr. H. Bito (University of Tokyo), then expanded in transformation-competent *E. coli* and purified with a Qiagen MaxiPrep kit, followed by extraction in a solution of phenol-chloroform-isoamyl alcohol (25:24:1) saturated with 10 mM Tris (pH 8.0) and 1 mM ethylenediaminetetraacetic acid (EDTA). The purified plasmid was packaged in an AAV8 serotype at the Boston Children’s Hospital vector core. For intersectional activity-dependent tagging, vectors were mixed in a 1:4 ratio of E-SARE-ERCreER to Cre_on_/Flp_on_-ChR2-eYFP or Cre_on_/Flp_on_-eYFP or Cre, respectively.

### Fiber optics

Fiber optic implants were constructed using ceramic ferrules (1.25 mm outer diameter; Thorlabs) with a 200 μm core, 0.39 numerical aperture multimodal fiber (Thorlabs). Fibers were cut to 4 mm length to target vCA1, and polished through a series of decreasing grit aluminum oxide polishing papers (Thorlabs). Fiber efficiency was measured with a light intensity meter (Thorlabs) used for calibrating light intensity during behavioral manipulations.

### Stereotaxic surgery

Stereotaxic viral infusion and optic fiber implantation surgery occurred 2-3 weeks before behavioral experimentation. Mice were anesthetized with vaporized 4% isofluorane in oxygen at 1.5 L min^-1^, head-fixed in a stereotaxic surgical frame (Stoelting), and maintained under anesthesia of 1-1.5% at 0.75 L min^-1^. Ophthalmic ointment was applied to the eyes to prevent drying. A skin incision was made to expose the skull. The skull was cleaned with 3% hydrogen peroxide and scored with a scalpel, and a craniotomy was drilled above the injection and fiber implant site. Viral vectors were injected bilaterally into vCA1 (from Bregma; anteroposterior (AP): -3.16 mm; mediolateral (ML): ± 3.5 mm; dorsoventral (DV): -3.5 and -4.1 mm) at a volume of 250 nL per injection at a rate of 100 nL min^-1^ using a motorized injector (World Precision Instruments) and glass syringe (Hamilton). The needle was left in the injection site for an additional 5 min after infusion before being slowly retracted. For optogenetic experiments, fiber optic cannulas were bilaterally implanted targeting vCA1 (AP: -3.16 mm; ML: ± 3.5 mm; DV: -3.5 mm) and were secured to the skull using C&B Metabond cement (Parkell) and dental cement (Lang). Mice were injected subcutaneously with banamine (2.5 mg kg^-1^) diluted in sterile saline (0.9%) to provide postoperative analgesia.

### Contextual Fear Conditioning

All mice were handled for 1-2 min daily for 3-4 days before behavioral testing. For optogenetic experiments, mice were then habituated to patch cords by allowing them to explore a novel cage while tethered to patch cables for 5 min. Mice were transported in their homecage from the vivarium to the holding room adjacent to the testing room at least 1 hour prior to experimentation. For experimental timelines involving optogenetic manipulation, mice were tethered to patch cables for all behavioral sessions regardless of whether laser light was delivered.

Behavioral testing was conducted in 30.5 (length) x 24 (width) x 21 (height) cm^3^ conditioning chambers (Med Associates) with aluminum side walls, a Plexiglass door and ceiling, and a vinyl back wall. Conditioning chambers were housed within a larger sound-attenuating chamber. An overhead light and fan delivering ∼65 dB ambient noise was present throughout every session. Two distinct contexts with unique sensory features were used – a conditioning context and alternate context. The conditioning context consisted of a stainless steel rod floor (36 rods spaced 8 mm apart measured center to center), and was cleaned and scented with 70% ethanol. The alternate context contained a curved plastic wall insert, a plastic floor covered with corncob bedding, and was cleaned and scented with isopropanol.

Contextual fear conditioning consisted of three 2 s 0.75 mA scrambled foot shocks delivered 180, 240, and 300 s after the start of the session. The mice were removed from the context 30 s after the last shock. Foot shocks were omitted for unconditioned mice. Baseline freezing is defined as the average freezing behavior during the first 3 min pre-shock period of this session.

Extinction training typically consisted of 5 daily sessions of 10 min unreinforced exposure to the conditioning context (exception: 5 min sessions for Fig. 2). The average freezing behavior from the first 3 min of each extinction session is reported, as context-evoked freezing is typically highest during this period^45^. Mice that were not extinguished remained in their home cage until the test session.

The reminder shock session consisted of a single 2 s 0.75 mA shock delivered 180 s after the start of the session in the conditioning context, and mice were removed 30 s later. Average freezing from the 3 min pre-shock period was reported as the ‘pre-reminder’ period.

The final test sessions prior to perfusion for c-Fos analysis consisted of 5 min exposure to the conditioned context.

All behavioral sessions were recorded at 30 frames per second using a near-infrared camera mounted to the interior door of the sound-attenuating chamber. The videos were manually scored for freezing behavior by an investigator blind to the experimental conditions. Freezing was defined as the absence of movement, with the exception of respiratory-related movements.

### Optogenetic manipulations during context tests

All optogenetic manipulation during context tests (fear retrieval, extinction retrieval, relapse, and alternate context) were 12 min in duration with light-on epochs occurring from min 0-3 and 6-9. NpHR-mediated photoinhibition was performed with a 568 nm laser (Opto Engine) delivered continuously with an intensity of 12-15 mW at the end of the fiber optic implant. ChR2-mediated photoexcitation was performed with a 473 nm laser (Opto Engine) delivered in 10ms pulses at 20 Hz with an intensity of 9-12 mW at the end of the fiber optic implant. All manipulations were performed bilaterally.

### Open Field Test

The apparatus was a 40 (length) x 40 (width) x 30.5 (height) cm^3^ box made of acrylic plastic with opaque walls. Overhead lights illuminated the center to ∼200 lux. Mice were connected to fiber optic patch cables and placed into the arena for 18 min and allowed to explore freely. Photostimulation consisted of 3 min light-on epochs from min 3-6, 9-12, and 15-18. The light parameters were identical to those used in contextual fear tests. The center zone was defined as an 18.5 x 18.5 cm^2^ zone in the center of the arena. Movement was recorded using a digital camera mounted above the arena at 15 frames per second and was analyzed with video-tracking software (ANY-maze).

### Appetitive Conditioning

All mice were food restricted to 80-90% of free-feeding weight beginning 5 days before behavioral testing and were weighed daily to maintain this deficit throughout the experiment. Mice were exposed to the rewarding solution (30% sucrose dissolved in water) for 3 consecutive days prior to testing delivered in a plastic weigh boat placed in their homecage. Appetitive conditioning was conducted in the same conditioning chamber and contextual features used for contextual fear conditioning, but with one region of the sidewall replaced with a sucrose reward port (Med Associates; ENV-303LPHD). The receptacle was connected via tubing to a syringe filled with the sucrose reward and loaded in a single speed syringe pump (PHM-200). Each reward was 20 μL in volume. Port entries were detected by breaks of an infrared beam incorporated into the port.

Contextual appetitive conditioning consisted of 30 unsignaled sucrose rewards. Beginning with the start of the trial, rewards were dispensed with an intertrial interval of 45 s + a variable duration with an average time of 15 s. Rewards were omitted after 6 (Fig. 6f-j) or 8 (Fig. 6a-e) days of conditioning for extinguished mice and were omitted entirely for unconditioned mice. Unconditioned mice received sucrose solution in their homecage following context exposure. The final test session prior to perfusion for c-Fos analysis was 5 min of context exposure with rewards omitted in all groups. For optogenetic manipulation of SST-INs following appetitive extinction, photoinhibition was delivered for the first 5 min of the 10 min test.

### Intersectional activity-dependent neural tagging

Activity-dependent tagging was induced with an intraperitoneal injection of 4-OHT (Sigma) at 10 mg kg^-1^ administered immediately following the behavioral experience of interest (e.g. fear conditioning or extinction retrieval). In Fig. 4a-e, mice were injected with vehicle if not administered 4-OHT. Following 4-OHT injections, mice remained in their homecage in a quiet room for multiple hours in order to minimize non-specific labeling. 4-OHT was first dissolved in dimethyl sulfoxide (DMSO) to a concentration of 40 mg mL^-1^, then diluted in a 1% Tween-80 solution in sterile saline (0.9%) to a final concentration of 1 mg mL^-1^.

### Tissue preparations and immunohistochemistry

For c-Fos analysis, perfusion began 90 min following context exposure. Mice were deeply anesthetized with isofluorane and transcardially perfused with chilled 1x phosphate buffered saline (PBS) followed by 4% paraformaldehyde (PFA) in 1x PBS. Brains were extracted and post-fixed overnight in 4% PFA at 4 °C and then transferred to 30% sucrose in 1x PBS for 2 days at 4 °C until saturated. 35 μm coronal sections were collected on a cryostat (Leica Microsystems) and stored in cryoprotectant at −20 °C.

For immunohistochemistry, floating sections were washed six times for five minutes in 1x PBS and blocked at room temperature for 1 h in 10% normal goat serum (NGS; Jackson ImmunoResearch; 005-000-121) in 1x PBS with 0.3% Tween-20 (PBST). Sections were incubated with primary antibodies - 1:2000 rabbit anti-cFos (Synaptic Systems; 226 003), 1:500 chicken anti-GFP (Invitrogen; A11122), or 1:1000 rabbit anti-SST (Thermo Fischer Scientific; PA5-82678) - diluted in 5% NGS in 1x PBST overnight at 4 °C. Sections were rinsed three times in 1x PBS and incubated in secondary antibodies of 1:500 goat anti-rabbit Alexa Fluor 647 (Jackson ImmunoResearch; 111-605-144) or 1:500 donkey anti-chicken Alexa Fluor 488 (Jackson ImmunoResearch; 703-545-155) in 1x PBST for 2 h at room temperature. Sections were rinsed three times in 1x PBS, mounted onto slides, and coverslipped with ProLong Gold antifade medium containing 4,6-diamidino-2-phenylindole (DAPI; Invitrogen; P36931). All incubations and washes done with slight agitation on an orbital shaker.

### Imaging and cell count quantification

Fluorescent confocal images of immunolabeled tissue were acquired on a Zeiss LSM 780 with a 20x objective and suitable filter sets in tiled Z-stack images (4 μm steps) that were stitched using the Zen Black software (Carl Zeiss). For each experiment, images were acquired with identical laser intensity, gain, and pinhole settings. Immunopositive cell bodies were counted manually in ImageJ (NIH) from contours of regions of interest by an experimenter blind to the experimental conditions. Area measurements derived from ImageJ contours was used to calculate estimate of cell count density. To obtain DAPI^+^ density estimates used in Supplementary Fig. 2, DAPI^+^ cells were quantified from a selection of 8 sample regions from vCA1 s.p. and s.o. Finally, the mean DAPI^+^ density estimate from these regions was multiplied by the total contour area to determine total estimated DAPI^+^ density for each animal.

### Electrophysiology

Mice were anesthetized with isoflurane before decapitation. Brains were extracted and submerged in ice-cold, carbogen-bubbled (95% oxygen, 5%CO_2_) sucrose cutting solution containing (in mM): 210.3 sucrose, 26.2 NaHCO_3_, 11 glucose, 2.5 KCl, 1 NaH_2_PO_4_, 0.5 sodium ascorbate, 4 MgCl_2_, and 0.5 CaCl_2_. Horizontal slices containing vCA1 were prepared on a VT1200S Vibratome (Leica Microsystems). Slices were recovered for 40 min at 35 °C in carbogen-bubbled artificial cerebrospinal fluid (ACSF) containing (in mM): 119 NaCl, 26.2 NaHCO_3_, 11 glucose, 2.5 KCl, 1 NaH_2_PO_4_, 2 MgCl_2_, and 2 CaCl_2_. Slices were subsequently maintained and recordings were performed at room temperature. For voltage-clamp recordings, whole-cell electrodes (4-6 MΩ) were pulled from borosilicate glass and filled with a low-chloride internal solution containing (in mM): 120 cesium methanesulfonate, 10 HEPES, 10 sodium phosphocreatine, 8 NaCl, 1 QX-314, 0.5 EGTA, 4 Mg-ATP and 0.4 Na-GTP. The internal solution was adjusted to pH 7.22 and 290–300 mOsm. Slices were visualized on an upright differential interference contrast microscope, and LED-coupled (Prizmatix) 40x objectives were used to identify fluorescently labeled cells. SST-INs were identified based on tdTomato fluorescence with high membrane resistance (>100 MΩ) or low capacitance (<90 pF) as confirmation of interneuron identity.

Spontaneous EPSCs and IPSCs were isolated by clamping the neurons at -60 or 0 mV, respectively. A total trace duration of at least 5 min was sampled at each potential for spontaneous currents. Data was low-pass filtered at 10 kHz and acquired at 10 kHz using Multiclamp 700B Microelectrode Amplifier (Molecular Devices) and pClamp 10 software (Molecular Devices, version 10.3.1). Cells exhibiting low health (>100 pA holding current), access resistance changes of >20%, or unstable synaptic responses were excluded from the analysis. All analyses were performed using Easy Electrophysiology.

### Statistics

All statistical tests and corresponding results and significance are indicated in the figure legends. Statistical analyses and figure generation was performed in Prism 10.10 (GraphPad Software). All data expressed as mean ± s.e.m. The α value was set at 0.05 for all analyses.

