## Supplementary Material for "Ventral hippocampal interneurons govern extinction and relapse of contextual associations"

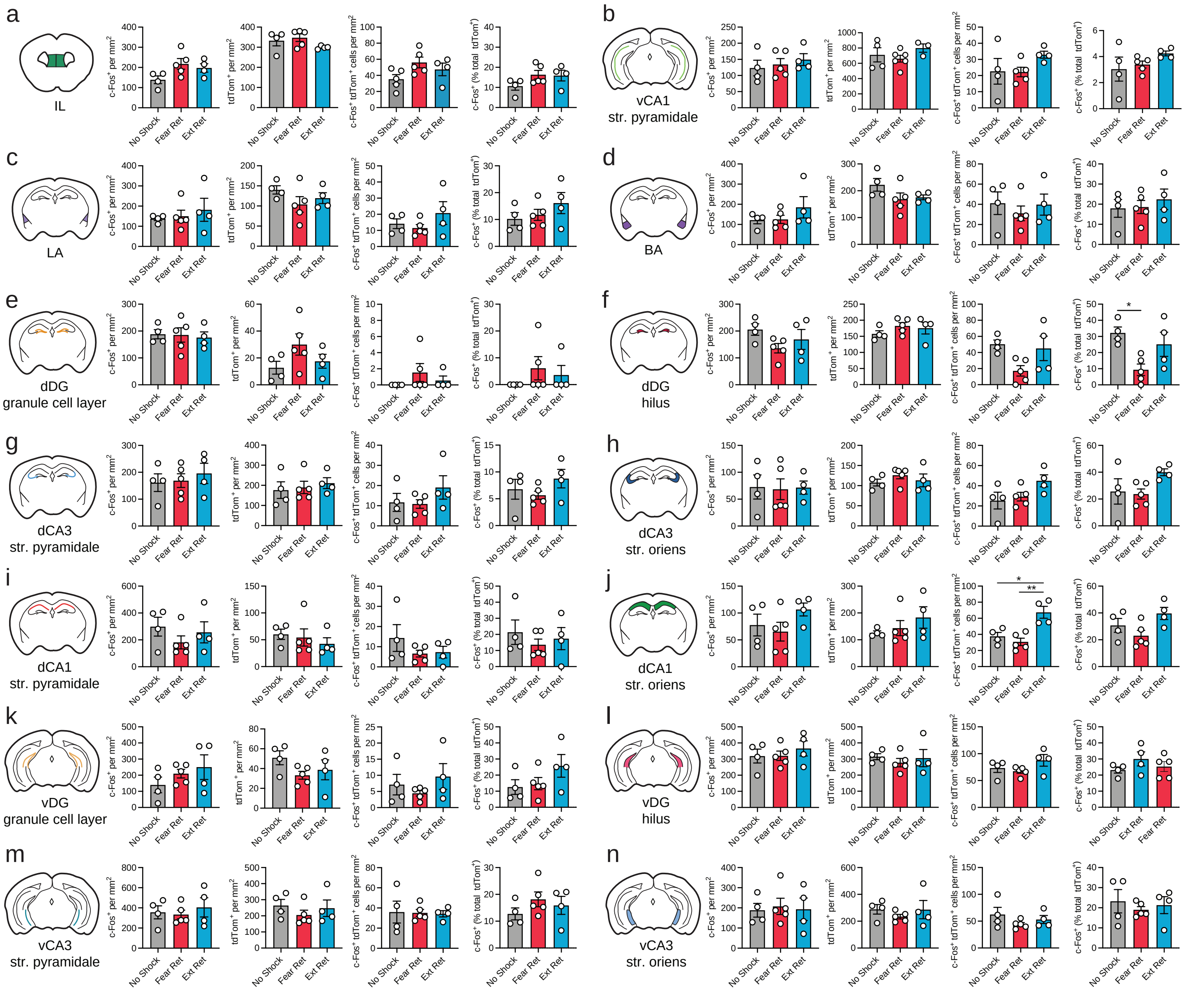


**Supplementary Figure 1. Additional regions of interest for c-Fos-based analysis of general and SST-IN-specific following fear or extinction retrieval, related to Fig. 1. (a-n)** Quantification of c-Fos+ cell density, tdTom+ cell density, c-Fos+ tdTom+ cell density, and c-Fos+ (% total tdTom+ cells) from **(a)** infralimbic medial prefrontal cortex (IL), **(b)** vCA1 str. pyramidale, **(c)** lateral amygdala (LA), **(d)** basal amygdala (BA), **(e)** dorsal dentate gyrus (dDG) granule cell layer, **(f)** dDG hilus, **(g)** dorsal CA3 str. pyramidale, (**h**) dorsal CA3 str. oriens, **(i)** dCA1 str. pyramidale, **(j)** dCA1 str. oriens, **(k)** vDG granule cell layer, **(l)** vDG hilus, **(m)** vCA3 str. pyramidale and (n) vCA3 str. oriens related to Fig. **1a-f** analyzed by one-way ANOVA. c-Fos+ (% total tdTom+ cells) from dDG hilus: $F_{(2, 10)} = 5.72$, $p < 0.05$ . c-Fos+ tdTom+ cell density from dCA1 str. oriens: $F_{(2, 10)} = 12.0$, $p < 0.01$. No Shock: $n = 4$; Fear Ret: $n = 5$; Ext Ret: $n = 4$. Data are presented as mean ± s.e.m. **(m)** lateral amygdala. **(n)** basal amygdala. * $p < 0.05$, ** - $p < 0.01$ Tukey's post-hoc test.

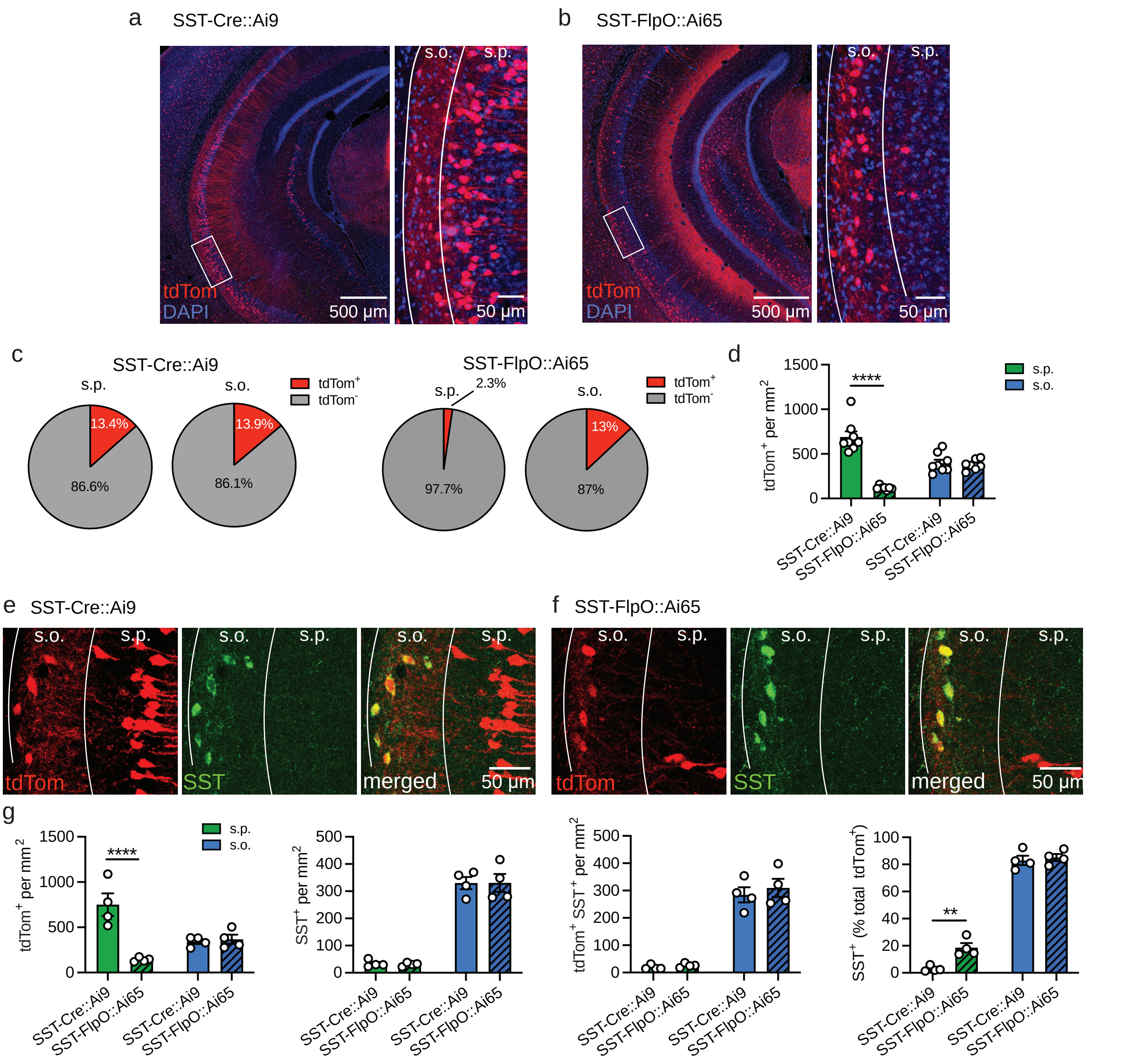


**Supplementary Figure 2. Comparison of recombination selectivity in SST-Cre versus SST-FlpO mice, related to Fig. 1. (a-b)**: Representative image of vCA1 in **(a)** SST-Cre::Ai9 and **(b)** SST-FlpO::Ai65 mice. Solid white box indicates region of magnification in right panel. **(c)** tdTom$^+$ and tdTom$^-$ cells as a proportion of DAPI$^+$ cells in vCA1 s.p. and s.o. layers. **(d)** tdTom$^+$ cell density in vCA1 s.p. and s.o. layers analyzed by two-way ANOVA. $F_{(1, 26)} = 45.2$, $p < 0.0001$. **(e-f)** Representative image of SST immunolabeling and tdTom expression in vCA1 of **(e)** SST-Cre::Ai9 and **(f)** SST-FlpO::Ai65 mice. **(g)** tdTom$^+$, SST$^+$, tdTom$^+$ SST$^+$ cell density, and SST$^+$ cells (% of tdTom$^+$ cells) in vCA1 s.p. and s.o., analyzed by two-way ANOVA. tdTom$^+$ density: $F_{(1, 12)} =$ 21.4, $p < 0.001$. SST$^+$ (% total tdTom$^+$): $F_{(1, 12)} = 5.12$, $p < 0.05$. **a-d:** SST-Cre::Ai9: $n = 8$; SST-FlpO::Ai65: $n =$ 7. **e-g**: SST-Cre::Ai9: $n = 4$; SST-FlpO::Ai65: $n = 4$. Data are presented as mean ± s.e.m. ** - $p < 0$ .001, **** - $p < 0.0001$ by Šidák's post hoc test.

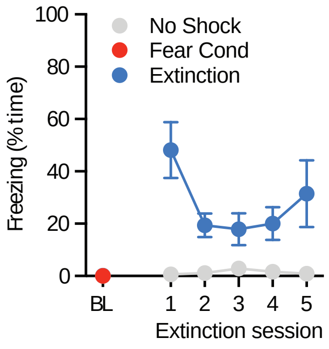


**Supplementary Figure 3. Freezing levels for mice used in synaptic electrophysiology experiments, related to Fig. 2. (a)** Freezing was quantified during the pre-shock baseline period during contextual fear conditioning and during each 5 min extinction session. No Shock: $n$ = 6; Fear Cond: $n$ = 4; Extinction: $n$ = 5. Data are presented as mean ± s.e.m.

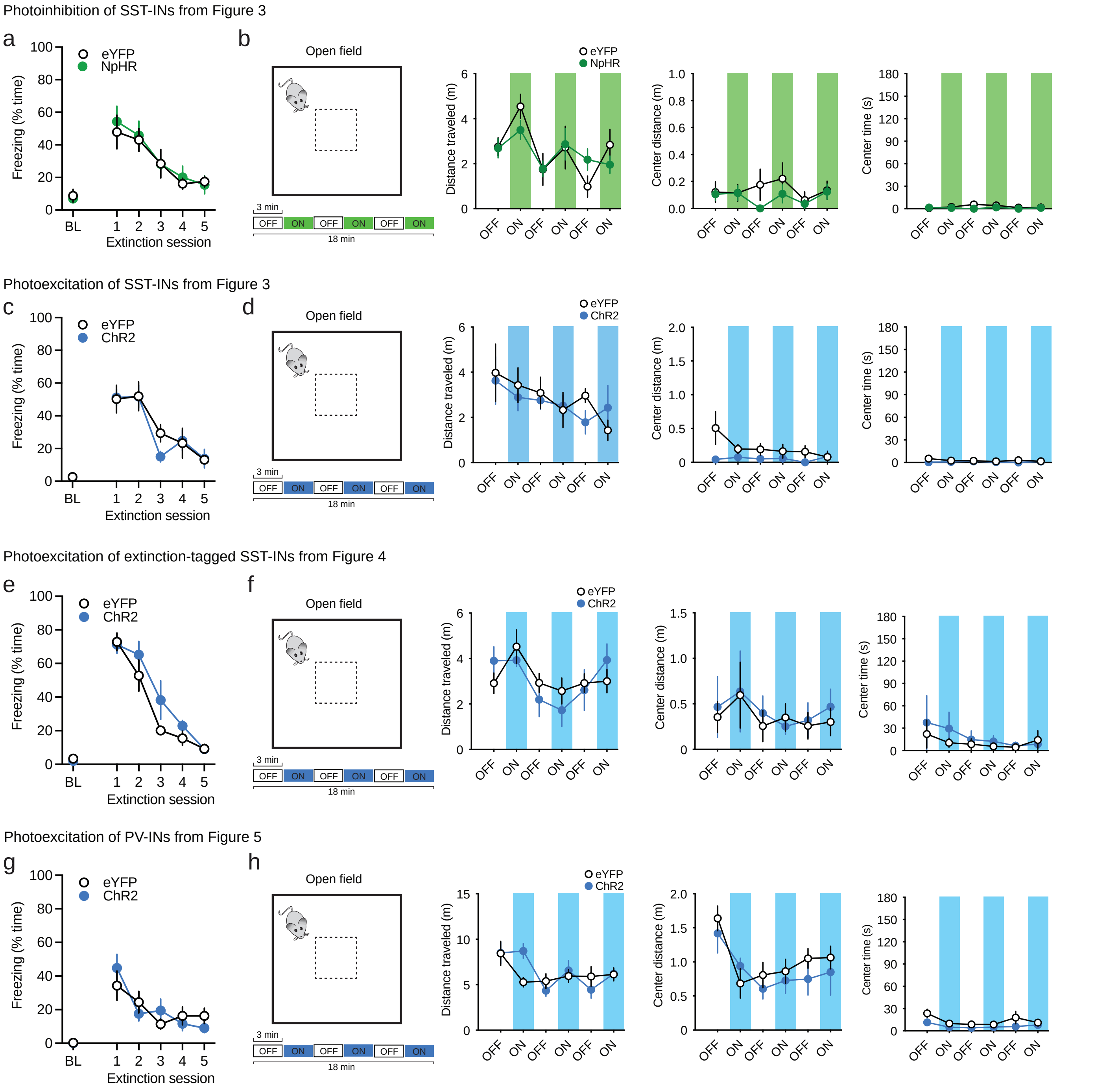


**Supplementary Figure 4. Extinction-related freezing and open field metrics for optogenetic manipulations of SST-INs, extinction-related SST-INs and PV-INs, related to Figs. 3-5. (a-b)** Additional analysis associated with photoinhibition of SST-INs in Fig. 3. of **(a)** freezing during the baseline period of the training session and first 3 min of each of the 5 extinction sessions, and **(b)** total distance traveled, center distance, and time in center from open field test. Photostimulation occurred in 3 min epochs beginning 3, 9, and 15 min into the 18 min test with light parameters identical to manipulations in fear conditioning contexts. **(c-d)** Analysis equivalent to that shown in **a-b** but associated with photoexcitation of SST-INs in Fig. 3. **(e-f)** Analysis equivalent to that shown in **a-b** but associated with photoexcitation of extinction-tagged SST-INs in Fig.4. **(g-h)** Analysis equivalent to that shown in **a-b** but associated with photoexcitation of PV-INs in Fig. 4. **a-b**: eYFP: $n$ = 6; NpHR: $n$ = 6. **c-d**: eYFP: $n$ = 5; ChR2: $n$ = 4. **e-f**: eYFP: $n$ = 5; ChR2: $n$ = 4. **g-h**: eYFP: $n$ = 6; ChR2: $n$ = 6. Data are presented as mean ± s.e.m.

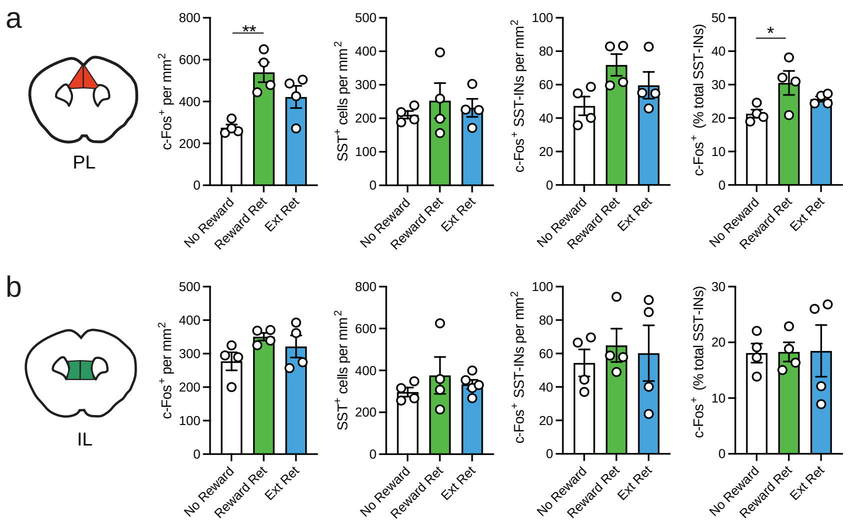


**Supplementary Figure 5. Additional analysis of SST-IN activation during retrieval of reward extinction, related to Fig. 6. (a-b)** c-Fos⁺ cell density, SST-IN density, c-Fos⁺ SST-IN density, and c-Fos⁺ SST-INs (% total SST-INs) from (**a**) prelimbic cortex (PL) and (**b**) infralimbic cortex (IL) during retrieval of an extinguished contextual reward memory analyzed by one-way ANOVA. c-Fos⁺ cells per $mm^2$: $F_{(2, 9)} = 10.0$, $p < 0.01$. c-Fos+ (% total SST-INs): $F_{(2, 9)} = 4.28$, $p < 0.05$. No Reward: $n = 4$; Reward: $n = 4$; Extinction: $n = 4$. Data are presented as mean ± s.e.m. * - $p < 0.05$; ** - $p < 0.01$ by Tukey's post-hoc test.
